# Start&Stop - a PhysiCell and PhysiBoSS 2.0 add-on for interactive simulation control

**DOI:** 10.1101/2024.12.13.628298

**Authors:** Riccardo Smeriglio, Roberta Bardini, Alessandro Savino, Stefano Di Carlo

## Abstract

In computational biology, *in silico* simulators are vital for exploring and understanding the behavior of complex biological systems. Hybrid multi-level simulators, such as `PhysiCell` and `PhysiBoSS 2.0`, integrate multiple layers of biological complexity, providing deeper insights into emergent patterns. However, one key limitation of these simulators is the inability to adjust simulation parameters once the simulation has started, which hinders the interactive exploration and adaptation of dynamic protocols ranging from biofabrication to *in vitro* pharmacological testing. To address this challenge, we introduce the `Start & Stop` add-on for `PhysiCell` and `PhysiBoSS 2.0`. This add-on offers multi-level state preservation and multi-modal stop control, triggered by simulation time or cell conditions, enabling users to pause a simulation, adjust parameters, and then resume from the exact halted state. We validate `Start & Stop` using two well-established `PhysiBoSS 2.0` use cases, a tumor spheroid 3T3 mouse fibroblasts use case under tumor necrosis factor (TNF) stimulation, and a lung cancer cell line invasion simulation, demonstrating that it preserves the simulator’s original behavior while enabling interactive configuration changes that facilitate the exploration of diverse and adaptive treatment strategies. By enhancing flexibility and user interaction, `Start & Stop` makes `PhysiCell` and `PhysiBoSS 2.0` more akin to real *in vitro* scenarios, thus expanding the range of potential simulations and advancing more effective protocol development in a variety of applications.

## 1 Background

In computational biology, *in silico* simulators are vital for exploring and understanding the behavior of complex biological systems [1].

Hybrid multi-level simulators integrate multiple layers of biological complexity, providing deeper insights into emergent patterns involving biological properties and behaviors at different scales. Moreover, they enable the examination of population dynamics in various environments and under different conditions [3][11]. Among the many available multi-level simulators, `PhysiCell` and `PhysiBoSS 2.0` are two notable examples [15][4]. In *in silico* pharmacological research, these simulators help users understand and test drug administration schemas more quickly and cost-effectively than traditional trial-and-error methods [16].

`PhysiCell` is a multicellular simulation framework utilizing various solvers, such as a diffusion and transport solver for simulating chemical microenvironment interactions with cells, a mechanical solver for simulating intercellular mechanical interactions, and additional components to simulate cellular processes like growth, division, cell differentiation, and death [9]. To efficiently simulate secretion, diffusion, uptake, and decay of multiple substrates in large three-dimensional (3D) microenvironments, `PhysiCell` employs the `BioFVM` solver [8]. `PhysiBoSS 2.0` extends `PhysiCell` by integrating it with `MaBoSS`, a continuous-time Markovian simulator that models intracellular signaling networks and regulatory processes within cells [19]. It provides a broad range of use cases, from drug treatment analysis of tumors to cell differentiation [15].

In the context of multi-scale simulators like `PhysiCell` and `PhysiBoSS 2.0`, the possibility of monitoring and controlling the simulation based on cell conditions is becoming crucial in several applications. Relevant examples include the development of efficient biofabrication protocols for regenerative medicine [6][10][2] and *in silico* pharmacological testing [20]. In the latter case, the ability to monitor and control the simulation to adjust the protocol settings after the simulation has started is crucial. For example, researchers could simulate the adjustment of treatments in response to observed cellular behaviors, making simulations more comparable to real *in vitro* experiments. Additionally, users could simulate biological processes such as tumor progression by dynamically introducing mutations, such as knocking out a critical gene involved in cell proliferation or migration.

Both `PhysiCell` and `PhysiBoSS 2.0` support easy customization of various parameters, such as simulation duration, set of administered stimuli, and initial position of cells. However, monitoring and controlling the simulation once it has started is impossible. To overcome this limitation, this paper introduces the `Start & Stop` add-on for `PhysiCell` and `PhysiBoSS 2.0`, providing:

- *Simulation breakpoints*: enabling the pause of the simulation based on the simulation time or predefined cell conditions.
- *Simulation state save and restore*: enabling to save snapshots of the system state after pausing a simulation and to restore them to restart the simulation from a previously saved state.

These new features enhance user control during the simulation and enable a more interactive and adaptive exploration of stimulation protocols.

## 2 Implementation

The `Start & Stop` add-on allows users to pause a simulation, modify the configuration parameters, and then restart it, preserving the simulation state. The add-on operates on all relevant model layers:

- **Cellular aggregate conformation**: an agent-based model of all cells, their phenotypes, their states, and their positions at the core of `PhysiCell`,
- **Intracellular states**: Only available if `PhysiBoSS 2.0` is used. Intracellular states are computed using Boolean models of the intracellular regulation network. The `MaBoSS` add-on embedded in `PhysiBoSS 2.0` handles this layer.
- **Microenvironment**: reaction-diffusion Physical Differential Equations (PDEs) that represent the 3D environment divided into 3D voxels. Each voxel is characterized by the concentration of all the relevant diffusing substances.

Figure 1 shows a schema of the integration into `PhysiCell` and `PhysiBoSS 2.0` of the `Start & Stop` add-on. It relies on three primary operations detailed in the following sections: *auto-stop* (Section 2.1), *current state saving* (Section 2.2), and *state restoring* (Section 2.3).

**Fig. 1.**
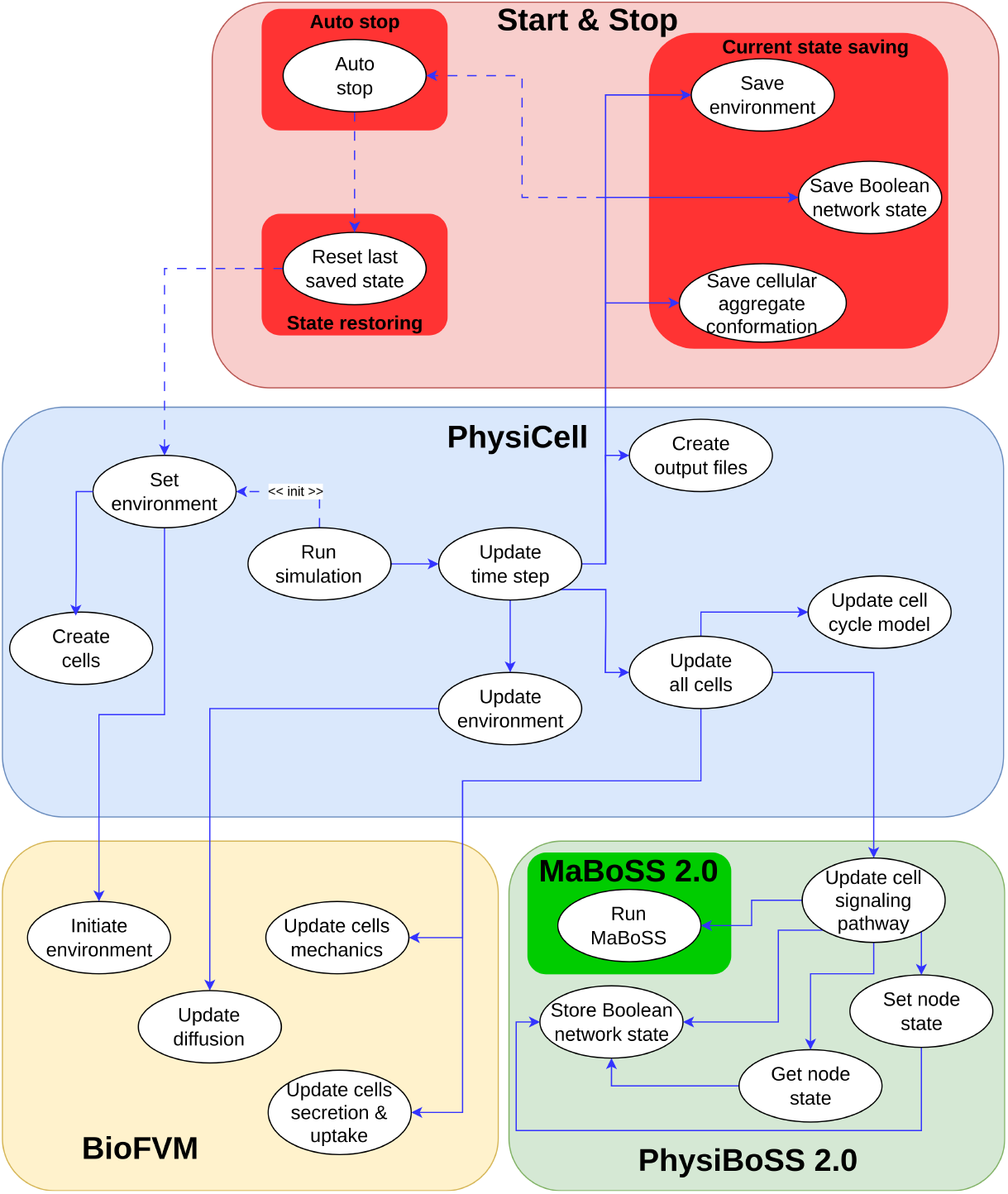
Integration of the Start & Stop add-on: This schema represents the integration of the Start & Stop add-on into the PhysiCell and PhysiBoSS 2.0 framework (adapted from [15]).

### 2.1 Auto-stop

Currently, in `PhysiCell` and `PhysiBoSS 2.0`, it is only possible to simulate a use case for a predefined amount of time, without the ability to monitor other parameters, unless the core simulator (i.e., `main.cpp` file) is modified.

Each use case in `PhysiCell` and `PhysiBoSS 2.0` natively includes the `custom.cpp` file, enabling `PhysiCell` and `PhysiBoSS 2.0` users to define custom functions. The `Start & Stop` add-on leverages this file to enable users to define a new function named `auto_stop` implementing a custom *monitoring system* and a set of *stop conditions*. An example of this function is provided in the `PhysiBoSS 2.0` sample use case `spheroid_tnf_model`, located in the file sample project intracellular/boolean/spheroid_tnf model_/custom_modules/custom.cpp.

With this functionality, we provide the possibility to stop the simulation by monitoring specific parameters instead of setting a predefined simulation.

### 2.1 Current state saving

To save the current simulation state, the `Start & Stop` add-on implements output streaming operators in the simulator’s core. Streaming operators are mechanisms offered by several programming languages that serialize and write complex data objects to external files, enabling the efficient storage of all simulation parameters. This ensures that essential information is preserved and can be reloaded when the simulation is resumed. These operators have been added in the `PhysiCell` classes such as `cells`, `cell container`, `custom`, `phenotype`, and `microenvironment`, as well as in `MaBoSS`. The `Start & Stop` add-on saves several files containing:

- Information about the *microenvironment*, with each voxel position and the concentration of all substances along with metadata;
- The state of the *cells*;
- The *position* (*x, y*, and *z* coordinates) of all the cells in the 3D environment;
- The `PhysiCell` global parameters, such as the current time, the next saving time, and other parameters;
- If using `PhysiBoSS 2.0`, each cell’s intracellular state is represented as the state of a Boolean model; if not, the intracellular state is not saved.

Importantly, all generated files are *human-readable*. Files are saved each time the simulation stops. Since the simulations can contain a huge number of cells and agents, in the case of multiple stops, the files are overwritten, thereby reducing storage requirements. The storage occupied by the saved files can vary as a function of the complexity of the simulation; complex simulations generate larger files. Examples of the storage requirements for the use cases presented in this paper are reported in section 3.3

### 2.3 State restoration

When a simulation is resumed using the `Start & Stop` add-on, the simulator reads all the state files saved at the last stop of the simulation. It restores all objects to the exact state they had when the simulation was interrupted, utilizing input streaming operators. This ensures that the multi-scale state of the entire simulator is fully restored before re-entering the main simulation loop. As a result, the simulation resumes in the exact state it was in when it was stopped. The integration in `PhysiCell` and `PhysiBoSS` is showed in Figure 1

### 2.4 Usage

To configure all the parameters for the simulation, `PhysiCell` and `PhysiBoSS 2.0` use an `XML` file. The `Start & Stop` add-on extends this file to contain two additional flags: `start_and_stop` and `auto_stop`. To enable the `auto_stop` functionality, the user must set the `auto_stop` flag to `true`. In this way, if the stopping conditions are met during the simulation, the simulation stops, and the simulation state is automatically saved. If the conditions are not met, the simulation continues until the pre-set time is reached. It is important to note that, even in the latter case, the simulation state is saved at the end of the simulation, allowing the user to resume it later. To restart a simulation from the last saved state, the user must set the `start_and_stop` flag to `true`. This ensures that the simulation resumes by loading the last saved state. An illustration of the functioning of the `Start & Stop` add-on in the context of a generic simulation is presented in Figure 2.

**Fig. 2.**
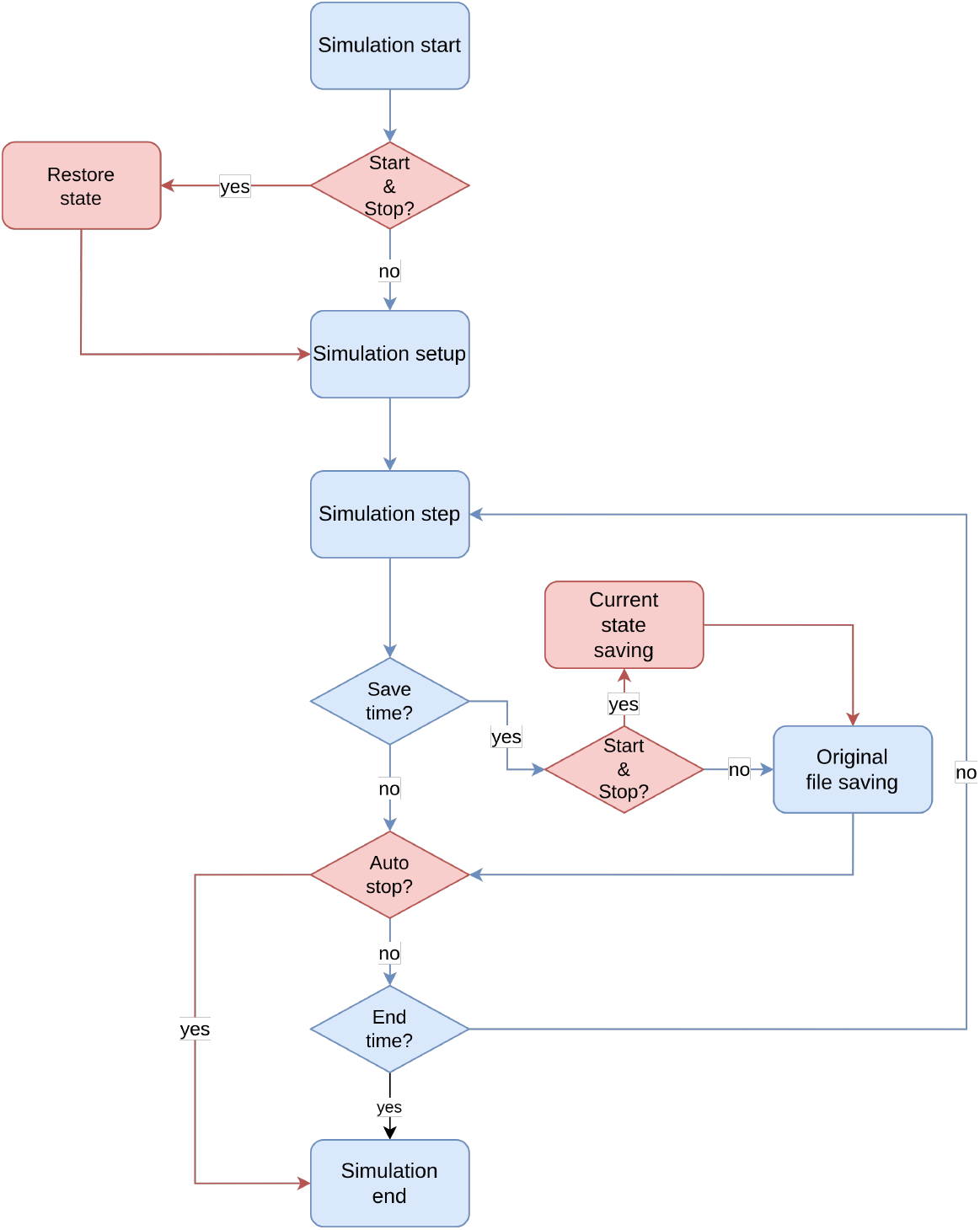
Start & Stop add-on functioning for a simulation. This flow chart illustrates a generic simulation in PhysiCell and PhysiBoSS 2.0, with the inclusion of the Start & Stop steps. The blue steps represent the native PhysiCell and PhysiBoSS 2.0 functioning, while the red ones represent the new Start & Stop add-on functionalities.

## 3 Results and Discussion

To demonstrate the functionalities of the `Start & Stop` add-on, we selected two wellestablished and validated `PhysiBoSS 2.0` use cases. One simulates a tumor spheroid 3T3 mouse fibroblasts under TNF stimulation, while the other simulates a lung cancer cell line invasion.

We selected these two use cases for their ability to support the verification and validation of the `Start & Stop` add-on and to showcase the added value of its new functionalities. In particular, the TNF cancer use case is a well-established and studied model [15]. Although it is not highly complex, featuring a Boolean model with 23 nodes, it includes the ability to select a stimulation protocol. This makes it an ideal test case for fully exploring the capabilities of the `Start & Stop` add-on, allowing us to evaluate its performance across different protocols and demonstrating its integration with `PhysiBoSS` by enabling protocol modifications during the simulation. Moreover, this use case serves as a valuable test not only for showcasing the capability of the `Start & Stop` add-on but also for verifying that it does not alter the core functionalities of the simulator. The model includes distinct phenotypes that are straightforward to analyze and track, particularly *apoptosis, necrosis*, and *survival* cell states. This characteristic facilitates the assessment of whether stopping and resuming the simulation preserves the expected biological behavior.

The second use case is more complex than the first, incorporating a Boolean model with 48 nodes. In addition to multiple cell cycle phases, this model simulates the transition from an *epithelial* to a *mesenchymal* phenotypes. This allows us to test the `Start & Stop` add-on in scenarios involving cell transitions, further validating its robustness. Although this use case does not include a predefined stimulation protocol, it still demonstrates the add-on’s flexibility by enabling the testing of different simulation conditions through modifications of internal cell parameters, such as triggering or inhibiting specific transitions.

This work presents four demonstrative tests to highlight both the seamless integration with `PhysiCell` and `PhysiBoSS 2.0` and the flexibility provided by the `Start & Stop` add-on. Specifically, we performed two tests, one for each use case, to demonstrate that the `Start & Stop` add-on does not introduce any bias into the simulators. Additionally, we conducted two demonstrations showcasing the practical use of the `Start & Stop` add-on. All tests related to the first use case are presented in Section 3.1, while those related to the second use case are presented in Section 3.2.

### 3.1 Use case I: TNF tumor spheroid

This use case examines the cellular dynamics of a tumor spheroid under various TNF regimes [12]. This multi-scale model focuses on 3T3 fibroblast spheroids to explore the complex dynamics observed during *in vitro* drug administration [13]. In particular, the simulation starts with the tumor spheroid composed of alive cells in a proliferative state. Subsequently, the tumor undergoes periodic TNF stimulations that possibly trigger them to transition from a proliferative state towards necrotic or apoptotic death states [15]. Intracellular signaling of the cells underlying the complex response to TNF stimulation relies on the Boolean model provided in [5]. When the system undergoes a too-intensive TNF administration, cells can develop resistance to TNF itself, escaping necrotic death and remaining in a stable state of survival (as described in [5], Figure 2). In this use case, the simulated TNF administration occurs in a pulsed manner, where TNF is delivered in pulses with a defined interval, referred to as the *period*. Pulse administration occurs at a fixed *concentration* and *duration*. Users can adjust these parameters for each single simulation; however, TNF stimuli have fixed *period, duration*, and *concentration* values along the entire simulation. For the tumor spheroid use case, different protocols, corresponding to different combinations of these parameters, yield varying outcomes, defining the cell state in terms of the number of *alive, necrotic*, and *apoptotic* cells (see Section 2.2). In this context, the `Start & Stop` add-on allows users to modify the stimulation protocol during the simulation.

The first step for the validation of the `Start & Stop` add-on is to demonstrate that it is not introducing differences in the functioning of the simulators. To verify this, one approach is to compare the outcomes of simulations using the same protocol, with and without the use of the `Start & Stop` add-on.

Since both `PhysiCell` and `PhysiBoSS 2.0` are stochastic simulators, the outcome of the same simulation can vary from one trial to another, making it challenging to determine if activating the `Start & Stop` feature introduces any differences. To address this, for the TNF tumor use case, we estimated the number of simulations needed to ensure the sample mean differs from the actual distribution mean by no more than *±*10 *alive* cells at the end of the simulation, using a method based on the Standard Error of the Mean (SEM) [17], which yielded 30 simulations. In particular, this method involves computing the SEM as the standard deviation of the sample divided by the square root of the sample size (*N*). Assuming the data follows an approximately normal distribution, the true mean is expected to lie within *±*1.96*× SEM* of the sample mean with 95% confidence. By setting this range to 10 cells and estimating the standard deviation from preliminary simulations, we solved for *N* in the equation 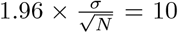 obtaining *N* = 30. To be more conservative, we considered 50 simulations.

As a drug administration scheme for testing, we performed two sets of 50 simulations using the same parameters: a TNF injection lasting 10 minutes every 150 minutes at a concentration of 0.005 TNF*/μm*^3^. In the first set, the simulations were executed continuously until completion without interruptions. In the second set, the simulations were stopped at every TNF administration period (every 150 minutes) and then resumed. This comparison allowed us to verify that introducing the `Start & Stop` does not alter the simulator’s behavior, demonstrating that the `Start & Stop` add-on does not introduce artifacts into the simulation process. Figure 3 reports the results. Figure 3-a, b, and c represent the box-plot of the distribution of *alive, necrotic*, and *apoptotic* cells, respectively, at each time step where the simulations with the `Start & Stop` add-on were stopped and restarted.

**Fig. 3.**
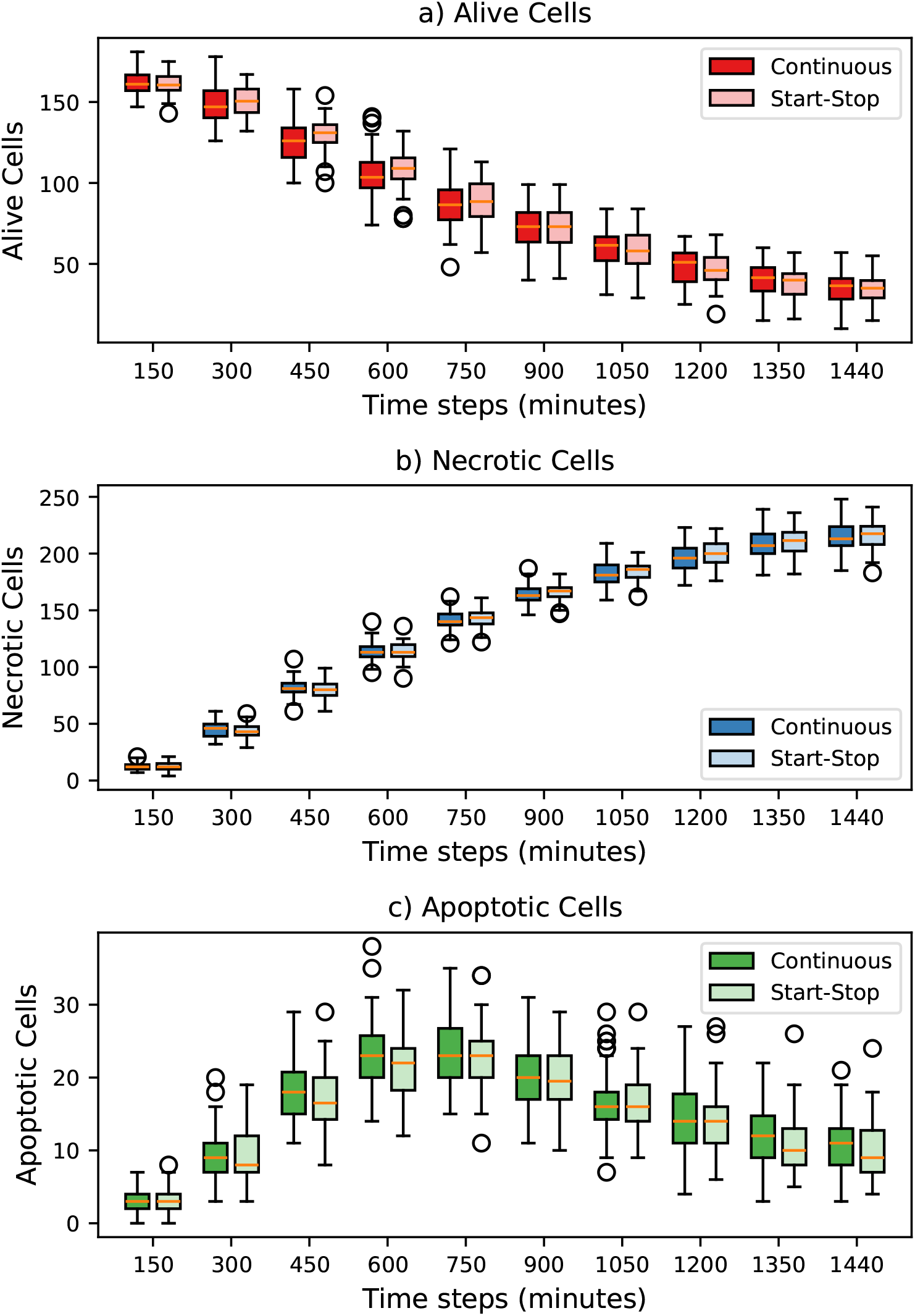
Simulation with and without the Start & Stop add-on: a) Comparison between simulations with and without activation of the Start & Stop add-on, for a simulation configuration with an administration every 150 minutes of 0.005 TNF*/μm*^3^ lasting 10 minutes, in particular for *alive* cells,b) *necrotic* cells, and c) *apoptotic* cells. Box plots are obtained by running 50 different simulations. In all simulations for the Start & Stop test, a stop and a restart occurred at each TNF administration time step.

Figure 3-a and Figure 3-b show that the behavior of both *alive, apoptotic*, and *necrotic* cells is consistent when comparing the simulations with and without the `Start & Stop` activation. In particular, *apoptotic* cells exhibit a higher standard deviation in their distribution, especially during the middle phase of the simulation (between 450 and 1050 minutes). This behavior is observed in both configurations, with and without the `Start & Stop` add-on, suggesting that it originates from the implementation of the original model itself.

To illustrate the practical relevance of the `Start & Stop` feature in *in vitro* drug administration protocol design, we present a specific use case. This feature is particularly useful for managing the risk of drug-induced resistance in tumor cells, a common undesirable outcome in pharmacological testing. In the case of 3T3 fibroblast tumor spheroids, tumor cells can survive treatment in two scenarios. First, when the drug stimulation is too weak to trigger necrosis. Second, when the cells develop resistance to the effect of the cytokines [5]. Both scenarios are modeled in this `PhysiBoSS 2.0` application.

Figure 4 illustrates both cases. The Figure shows a simulation with continuous TNF administration at a concentration of 0.1 TNF*/μm*^3^. At some point, almost all cells became resistant to the treatment, leading to the continuous growth of living cells, as shown in Figure 4-a. To identify resistant cells, we analyzed the state of their Boolean model and marked as resistant all stable states that resulted in survival, following the criteria described in [5].

**Fig. 4.**
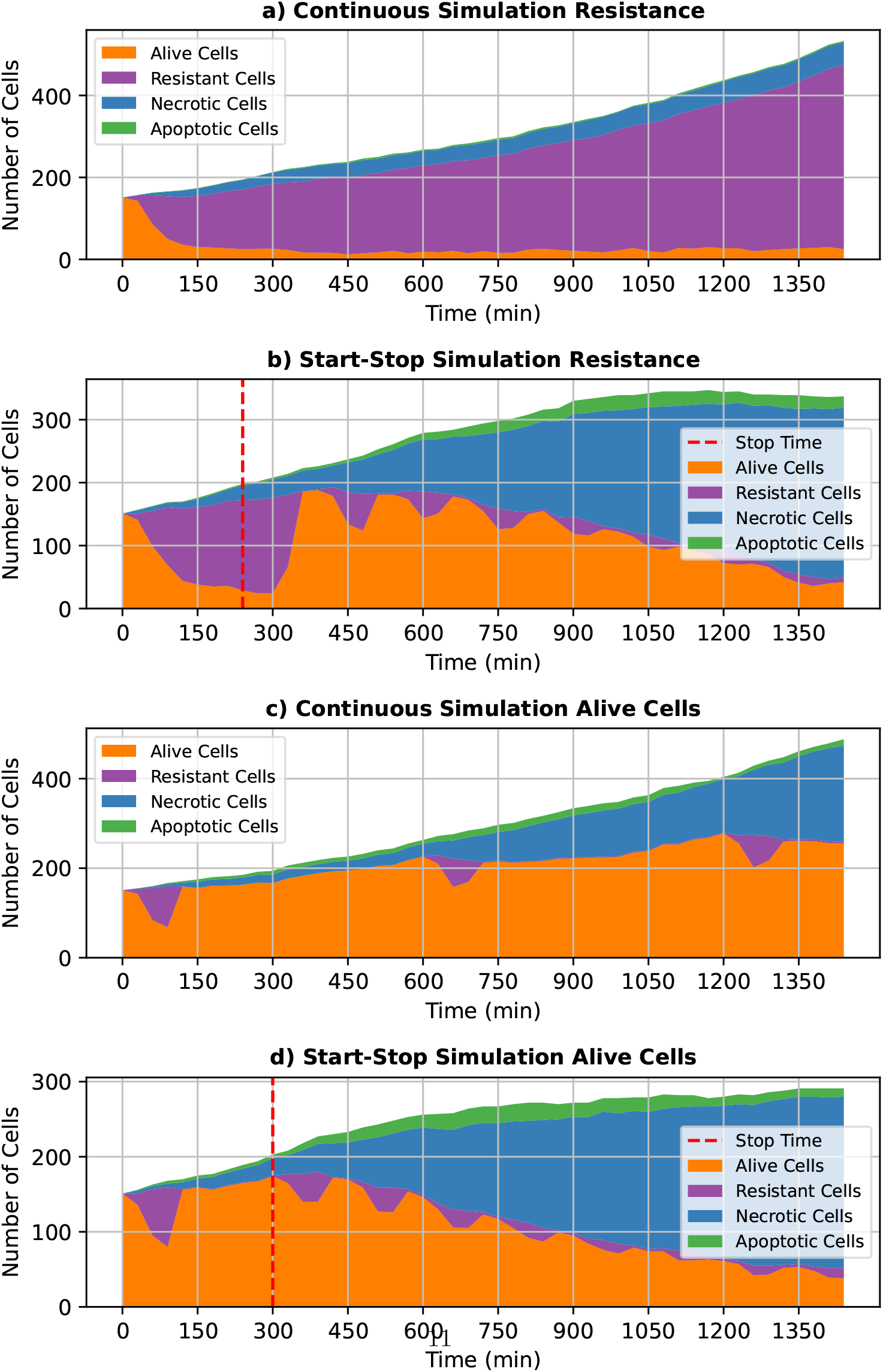
Application of the Start & Stop add-on in an in vitro drug administration simulation: a) Behavior of apoptotic, necrotic, alive, and resistant cells during the *in vitro* drug administration with continuous administration of 0.1 TNF*/μm*^3^ over time. Cells develop resistance to the therapy. b) Behavior of alive, apoptotic, necrotic, and resistant cells in a simulation starting similarly to Figure 4-a, but stopped at minute 240 and restarted with administration every 150 minutes of 0.005 TNF*/μm*^3^ for 10 minutes. c) Behavior of apoptotic, necrotic, alive, and resistant cells in a simulation with administration every 600 minutes of 0.005 TNF*/μm*^3^ for 10 minutes, where the *in vitro* drug administration efficacy is insufficient. d) Behavior of alive, apoptotic, necrotic, and resistant cells in a simulation starting similarly to Figure 4-c, but stopped at minute 300 and restarted with administration every 150 minutes of 0.005 TNF*/μm*^3^ for 10 minutes.

To demonstrate the capabilities of the `Start & Stop` add-on, we used the `auto-stop` function (see Section 2.1) to monitor and counteract cell resistance to TNF. Specifically, we created a monitor able to stop the simulation when more than 80% of the living cells acquire a resistant state. Figure 4-b shows that the simulation automatically stopped after 240 simulated minutes. At this point, we adjusted drug administration by providing stimulation every 150 minutes with 0.005 TNF*/μm*^3^ for 10 minutes. This change rapidly reduced cell resistance, allowing the drug to become effective.

Moreover, *in vitro* drug administration can fail due to insufficient stimulation. Figure 4-c illustrates this scenario, where we administered 0.005 TNF*/μm*^3^ every 600 minutes for 10 minutes. Although the *alive* cells did not develop resistance as in Figure 4-a, the treatment remained ineffective. To monitor this issue, we implemented another `auto-stop` function to track the number of *alive* cells and detect continuous tumor growth. As shown in Figure 4-d, the simulation stopped at simulated minute 300. We then modified the drug administration scheme to administrate every 150 minutes with 0.005 TNF*/μm*^3^ for 10 minutes. This adjustment successfully reduced the number of *alive* cells.

These two examples demonstrate the effectiveness and flexibility of the `Start & Stop` add-on.

### 3.2 Use case II: Cancer invasion

This use case describes a cancer cell line invasion model that includes different cell migration modes, with intracellular regulation governed by the Boolean model from [18]. This model builds on the Boolean network described in [7], which was validated using experimental data on transcriptome dynamics following TGFbeta–induced epithelial-to-mesenchymal transition (EMT) in lung cancer cell lines. It features two inputs: *ECMenv*, monitoring the Extracellular Matrix (ECM) status, and *DNA damage*, which accounts for DNA alterations triggering death signals. To also consider *Oxygen, GFs, TGFbeta*, and cell-cell contact, four additional inputs were introduced in [18]. The outputs, of the model include *CellCycleArrest, Apoptosis, EMT, ECM adh* (for cell adhesion), *ECM degrad* (for cell degradation), *Cell growth* (for tumor growth dynamics), and *Cell freeze* (for cell motility ability). The authors also included an EMT node to simulate the epithelial-to-mesenchymal transition [18]. As in use case I (see Section 3.1), for this second use case, we ran two sets of

50 simulations. This allowed us to support the comparison of the simulation states considering the stochasticity intrinsic to the simulators as performed in [18]. In the first set, simulations were run continuously for 2880 simulated minutes. In the second set, simulations were paused and resumed every 600 simulated minutes until reaching 2880 minutes in order to better capture the behavior of the involved cells. The results are shown in Figure 5. Figure 5-a shows the behavior of the *epithelial* cells, while Figure 5-b shows the behavior of *mesenchymal* cells. The distribution of both cells is consistent with and without the `Start & Stop` add-on with no significant differences highlighted. The results of this comparison again validate that the `Start & Stop` add-on does not alter the function of `PhysiCell` and `PhysiBoSS 2.0`.

**Fig. 5.**
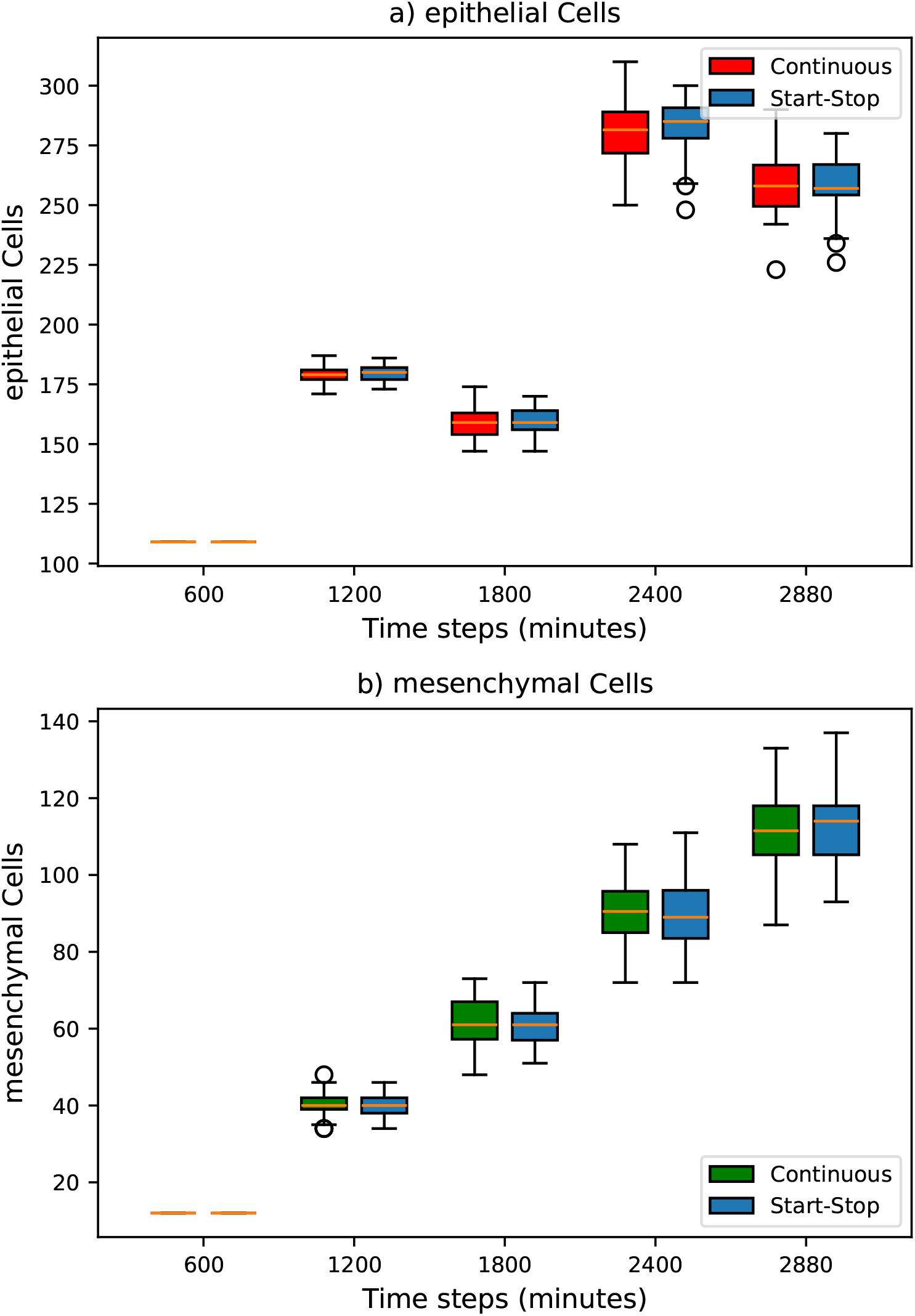
Simulation with and without the Start & Stop add-on: a) Comparison between simulations with and without activation of the Start & Stop add-on for *epithelial* cells. b) Comparison between simulations with and without activation of the Start & Stop add-on for *mesenchymal* cells.

As provided in [18], EMT is one of the key aspects underlying cancer invasion. This process can be influenced by intracellular factors, such as mutations, or external stimuli, including environmental changes, which may act as activators of specific disregulations that cause the cell to undergo EMT [18]. To demonstrate the application of the `Start & Stop` add-on in this context, we simulated the effect of intermittent blue light stimulation to induce the activation of the pro-invasive *SRC* protein, as described in [14]. Specifically, we modified the Boolean model presented in [18] (described in Section 3.2) by forcing the *SRC* node to remain active throughout the entire simulation. The results are presented in Figure 6.

**Fig. 6.**
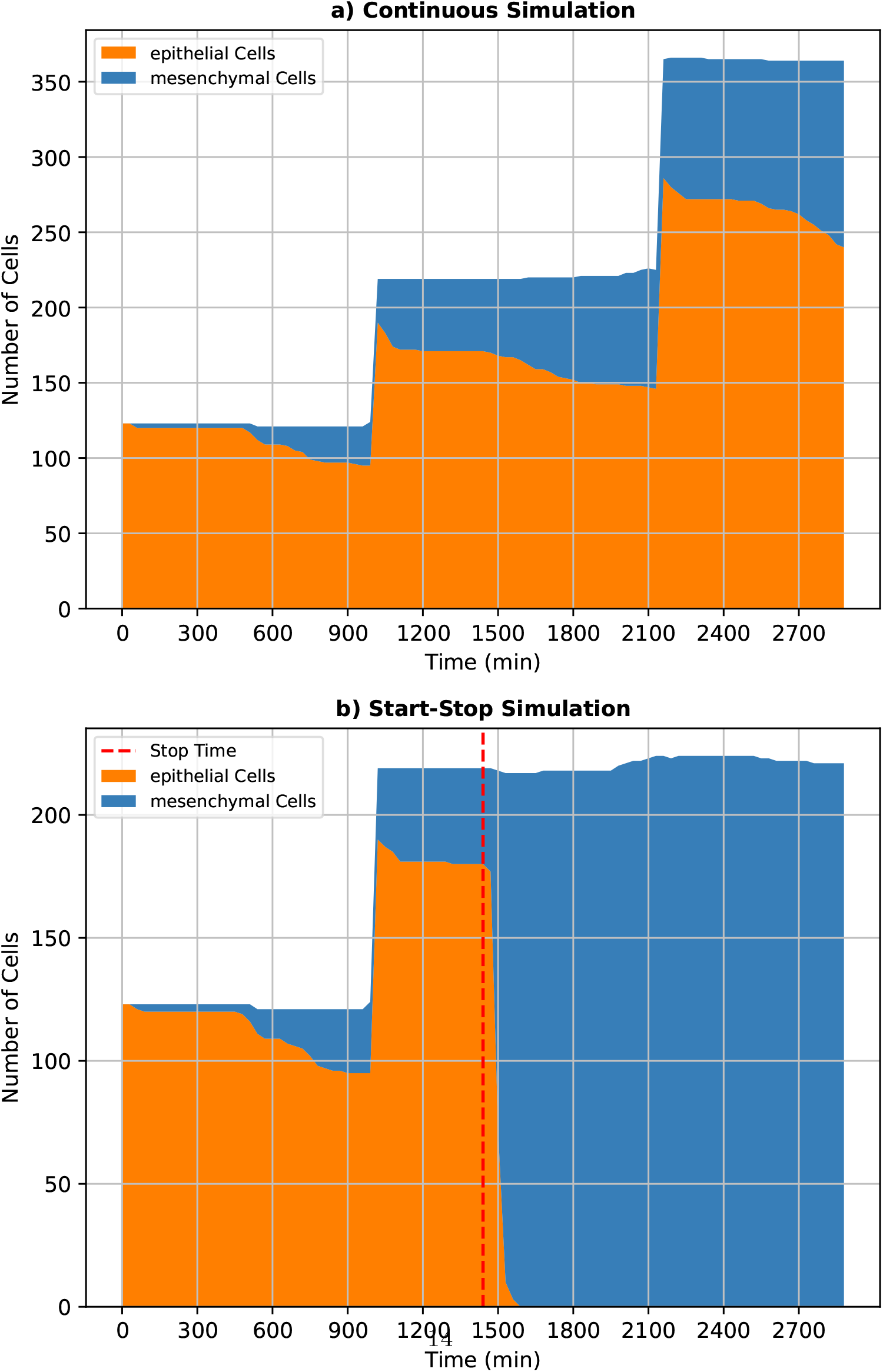
Application of the Start & Stop add-on in cancer invasion simulation: a) Behavior of *epithelial* and *mesenchymal* cells during the simulation. b) Behavior of *epithelial* and *mesenchymal* cells in a simulation that starts the same as Figure 6-a but is stopped at 1440 minutes and restarted with the *SRC* node active. In this second case, *epithelial* cells quickly transform into *mesenchymal* cells.

Figure 6-a illustrates the behavior of *epithelial* and *mesenchymal* cells under the dynamics of the *SRC* node throughout 2880 simulated minutes using the Boolean model presented in [18]. Initially, all cells are *epithelial*, but as the simulation progresses, they begin transitioning into *mesenchymal* cells. Figure 6-b shows the simulation described above during the first half (1440 simulated minutes) of the total 2880-minute simulation. At 1440 minutes, the simulation is paused, the `Start & Stop` add-on is activated, and during the state restoring phase, the intracellular Boolean model is replaced with the modified version as a strategy to model the optically-induced *SRC* activation. This triggers the disaggregation of cell junctions and thus, in turn, EMT [18]. As a result, in a few simulation steps, all the *epithelial* cells transition to a *mesenchymal* state.

### 3.3 Computational Costs

To further evaluate the impact of the `Start & Stop` add-on, we analyzed the storage and simulation time overhead introduced by the add-on compared to `PhysiCell` and `PhysiBoSS 2.0`.

Regarding the storage required to save the simulation state, it is proportional to the complexity of the model and simulation. In the first use case, at the end of a simulation with TNF administration every 150 minutes at a concentration of 0.005 TNF*/μm*^3^ for 10 minutes, the `Start & Stop` saved files occupied a total of 1.6 MB. Specifically, the simulation stored data for 263 cells, each with 23 nodes in the Boolean model. In the second use case, at the end of a simulation with the predefined initial parameters, the total storage occupied by the `Start & Stop` saved files was 3.2 MB. Since this use case is more complex than the previously discussed one, with a total of 377 cells and 48 nodes in the Boolean model, the occupied storage is twice as ample. In general, we can conclude that the overhead in terms of storage is not critical in the considered use cases but must be carefully considered in relation to the complexity of the model.

In terms of time overhead, the time of a `PhysiCell` or `PhysiBoSS` simulation including the `Start & Stop` add-on can be estimated as follows:

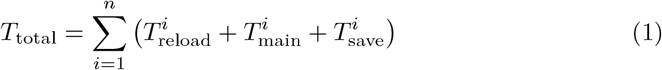

where *n* represents the number of simulation stops.

*T*_reload_ represents the simulation state reload time that depends on the number of cells and the dimension of the model. It can be computed as:

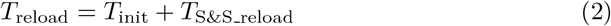

with *T*_init_ representing the time required to initialize the simulation, which also depends on the number of cells and the model size, and *T*_S&S reload_ representing the time needed to restore the last saved state. Please note that at the beginning of the simulation, *T*_reload_ = *T*_init_, since no previous state is reloaded.

*T*_main_ represents the time required for the execution of the main simulation loop.

Finally, *T*_save_ represents the time needed to save the files for the `Start & Stop` add-on.

The overhead caused by the `Start & Stop` add-on can be defined as:

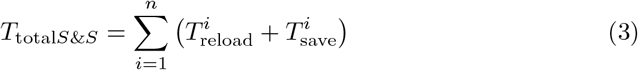

Figures 7 and 8 show the mean CPU time for each simulation phase as a function of the simulated time for the TNF cancer and cancer invasion use cases, respectively. These times were extracted from the same simulations presented in Figures 3 and 5. Figure 7 shows that *T*_main_ represents the most time-consuming phase of the simulation, and the overhead introduced to stop and resume the simulation is low (*T*_Reload_ and *T*_Save_). It is interesting to note that *T*_main_ gradually decreases as the simulation progresses. This reduction occurs because, as shown in Figure 3, an increasing number of cells are dying and they are skipped during updates, reducing the computational cost of the simulation. Additionally, the simulated time at the last stop is shorter than at the previous ones. Regarding the overhead introduced by the `Start & Stop` add-on, both *T*_save_ and *T*_reload_ increase as the total number of cells grows. Although the overhead increases, the most significant computational cost remains *T*_*main*_.

**Fig. 7.**
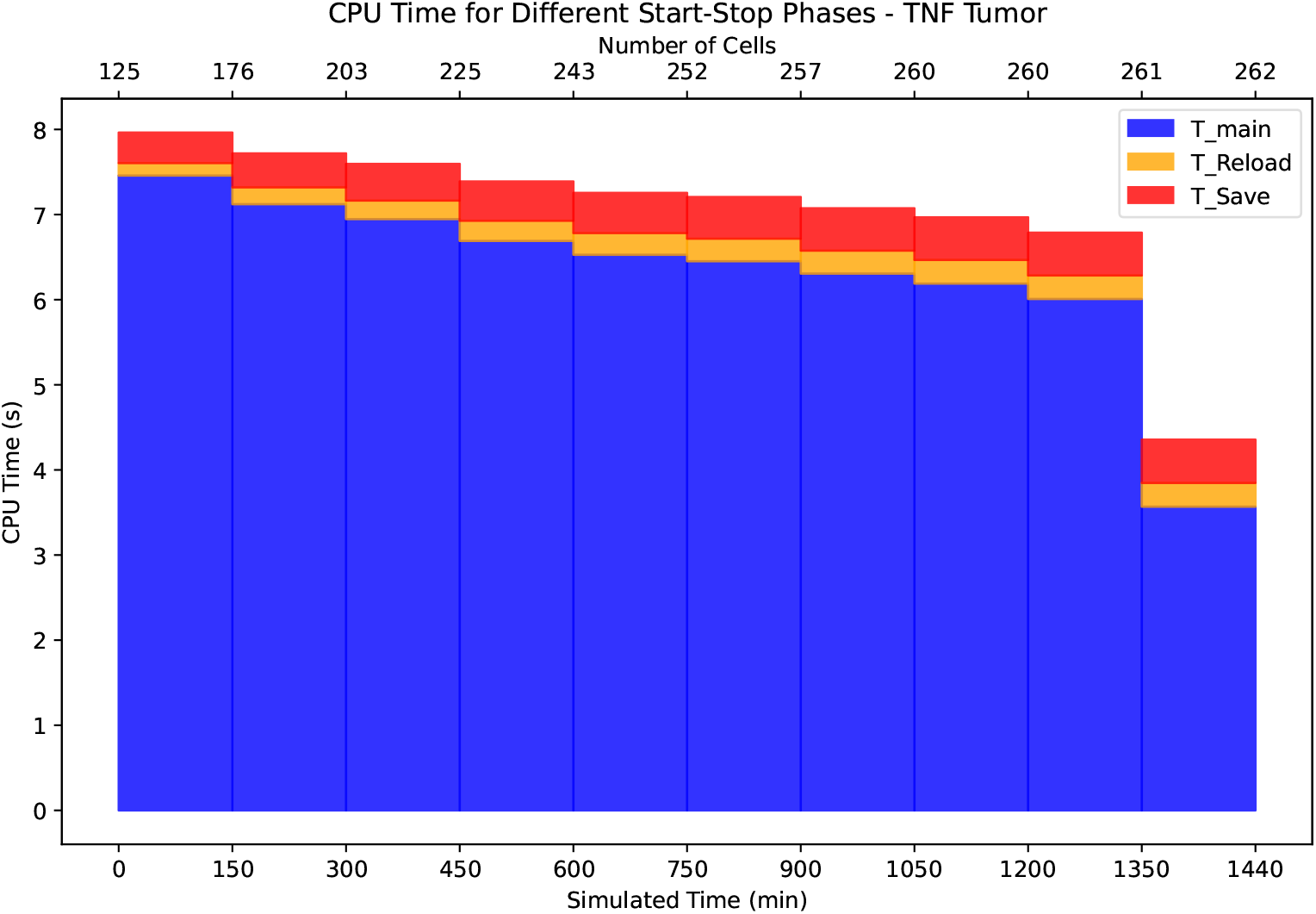
CPU time of the TNF cancer simulation as a function of the simulated time: This plot represents the CPU time required by the different phases of the simulation, shown as a function of both the simulated time and the number of cells at each stop for the TNF cancer use case.

**Fig. 8.**
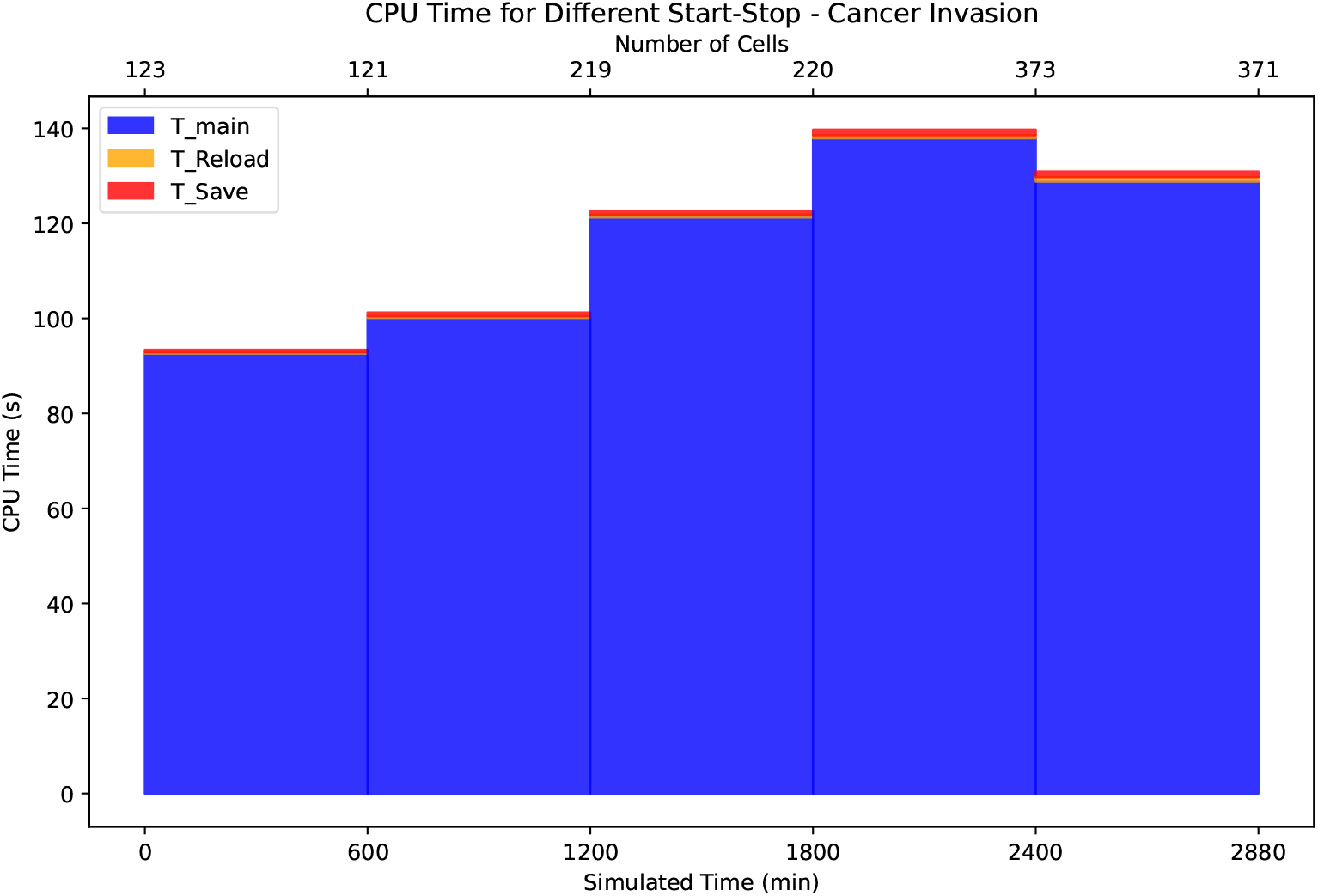
CPU time of the cancer invasion simulation as a function of the simulated time: This plot represents the CPU time required by the different phases of the simulation, shown as a function of both the simulated time and the number of cells at each stop for the cancer invasion use case.

This behavior is even more evident in the cancer cell line invasion use case, shown in Figure 8. Due to its greater complexity, this simulation requires significantly more CPU time for the execution of the main simulation loop, making the overhead introduced by the `Start & Stop` add-on almost negligible. However, as in the previous case, both *T*_save_ and *T*_reload_ increase as the total number of cells grows.

A final comparison between the mean total CPU time of the simulations with and without the `Start & Stop` add-on is shown in Table 1. From this table, it is evident that the inclusion of the `Start & Stop` add-on does not significantly increase the total CPU time of the simulation, as the values for continuous and `Start & Stop` simulations are comparable.

**Table 1.** Comparison of continuous and Start & Stop simulations for TNF Cancer and Cancer cell line Invasion.

|  | Time Continuous [s] | Time Start & Stop [s] |
| --- | --- | --- |
| **TNF Cancer** | $72.97 \pm 9.02$ | $71.90 \pm 3.43$ |
| **Cancer Invasion** | $581.06 \pm 16.47$ | $588.84 \pm 11.99$ |

## 4 Conclusions and Future Works

In this paper, we presented the `Start & Stop` add-on for `PhysiCell` and `PhysiBoSS 2.0`. This add-on enhances the flexibility of the simulators by:

- enabling the user to stop and restart the simulation from the same state at the point of interruption, as described in Section 2.3;
- allowing dynamic adjustments to a broad set of parameters, increasing user control and interaction with the simulation;
- facilitating the debugging of simulations in case of issues by allowing the user to monitor and easily interact with saved parameters in an accessible way.

The experiments described in Sections 3.1 and 3.2 demonstrated that the `Start & Stop` add-on does not alter simulation behavior when pausing and restarting. This functionality contributes to the computational ecosystem of hybrid and multi-level models in systems biology by enabling interactive and automated simulation control, thereby enhancing both representation and predictive power within the `PhysiCell` and `PhysiBoSS 2.0` framework.

Current limitations of this work include:

- As discussed in Section 2.2, saving and overwriting files at each stop minimizes storage usage but limits flexibility, as simulations can only be resumed from the last saved state. To overcome this, we plan to optimize storage and increase the frequency of saved states.
- Currently, the only ways to modify specific internal cell parameters, such as volume

or the state of the Boolean model, are either by manually editing the saved files, which is error-prone and may cause crashes, or by implementing a custom function to apply changes between the state-restoration phase and simulation restart. To improve this process, we plan to develop a user-friendly interface that simplifies parameter management.

In conclusion, the `Start & Stop` add-on can significantly expand the capabilities of the `PhysiCell` and `PhysiBoSS 2.0` simulator, making *in silico* experiments closer to *in vitro* experiments.

## 5 Availability and requirements

### 5.1 Project name

PhysiS&S

### 5.2 Project home page

https://github.com/smilies-polito/PhysiSandS

### 5.3 Operating systems

The project was developed and tested on Linux Ubuntu 22.04.4 LTS (GNU/Linux 5.15.0-131-generic x86 64)

### 5.4 Programming languages

`C++` for the add-on implementation.

`Python` for the test implementation.

### 5.5 Other requirements

All the dependencies to run the code independently are listed in the https://github.com/smilies-polito/PhysiSandSRepository.

### 5.6 License

BSD 3-Clause License.

### 5.7 Any restrictions to use by non-academics

Not applicable.

## Declarations

### Ethics approval and consent to participate

Not applicable.

### Consent for publication

Not Applicable.

### Availability of data and materials

The set of functions required for the `Start & Stop` functionality is implemented in `C++` and contained in the `add-ons/start and stop` folder, as required by `PhysiCell`. These functions need to be called in the `main.cpp` file of the selected use case. Implemented at the core of `PhysiCell`, it is possible to use the `Start & Stop` add-on for all `PhysiCell` and `PhysiBoSS 2.0` use cases. The full implementation and the script for reproducing the tests provided in this paper are publicly available on GitHub: https://github.com/smilies-polito/PhysiSandS. The add-on code is currently under review by the authors of the `PhysiCell` simulator as part of a pull request to the `PhysiCell` original repository: https://github.com/MathCancer/PhysiCell.

### Authors’ contributions

RS developed the code. RS and RB wrote the manuscript. RB, AS, and SDC supervised the work and reviewed the manuscript.

## Acknowledgment

We would like to thank Vincent Nöel, Miguel Ponce de Leon, Arnau Montagud, and the rest of `PhysiCell` and `PhysiBoSS 2.0` teams for their insightful suggestions and feedback.

